# The Crystal Structure of Bromide-Bound *Gt*ACR1 Reveals a Pre-Activated State in the Transmembrane Anion Tunnel

**DOI:** 10.1101/2020.12.31.424927

**Authors:** Hai Li, Chia-Ying Huang, Elena G. Govorunova, Oleg A. Sineshchekov, Meitian Wang, Lei Zheng, John L. Spudich

## Abstract

The crystal structure of the light-gated anion channel *Gt*ACR1 reported in our previous research article (*Li et al., 2019*) revealed a continuous tunnel traversing the protein from extracellular to intracellular pores. We proposed the tunnel as the conductance channel closed by three constrictions: C1 in the extracellular half, mid-membrane C2 containing the photoactive site, and C3 on the cytoplasmic side. Reported here, the crystal structure of bromide-bound *Gt*ACR1 reveals structural changes that relax the C1 and C3 constrictions, including a novel salt-bridge switch mechanism involving C1 and the photoactive site. These findings indicate that substrate binding induces a transition from an inactivated state to a pre-activated state in the dark that facilitates channel opening by reducing free energy in the tunnel constrictions. The results provide direct evidence that the tunnel is the closed form of the channel of *Gt*ACR1 and shed light on the light-gated channel activation mechanism.

**Impact Statement:** Substrate-induced structural changes in *Gt*ACR1 provide new insight into the chemical mechanism of natural light-gated anion conductance, and facilitate its optimization for photoinhibition of neuron firing in optogenetics.

## Introduction

*Gt*ACR1 is a light-gated anion channel discovered in 2015 (*Govorunova et al., 2015*) now widely used in optogenetics as a neuron-silencing tool. *Gt*ACR1 conducts both bromide and chloride ions effectively with higher relative permeability for the former substrate (*Govorunova et al., 2015*). We and the group of Karl Deisseroth independently determined X-ray crystal structures of the dark (closed) form of *Gt*ACR1 homodimer at 2.9 Å (*Kim et al., 2018; Li et al., 2019*). We proposed that the conductance pathway was attributable to a full-length intramolecular tunnel traversing each protomer from the extracellular side to the intracellular side of the membrane lined by mostly hydrophobic residues (*Li et al., 2019*). Current rectification by charges introduced inside but not outside the tunnel support our hypothesis that the tunnel serves as the anion-conducting path upon photoactivation (*Sineshchekov et al., 2019*). However, no substrate was found in either structure (*Kim et al., 2018; Li et al., 2019; Li et al., 2019*) despite the presence of chloride in the crystallization conditions. The mechanism of anion conductance is still elusive.

In the apo form (i.e. without anion substrate) (*Li et al., 2019*) the tunnel is narrowed by 3 constrictions blocking ion permeation: C1 in the extracellular half, C2 mid-membrane consisting of the photoactive retinylidene Schiff base and interacting residues, and C3 in the cytoplasmic half. In our model, retinal photoisomerization at C2 needs to open all three of these gates to form a conductive anion channel through the protein. However, structural changes at the two constrictions, C1 and C3, which are located on each side of the Schiff base, and their roles in the channel gating mechanism are unclear.

To address these questions, here we report the crystal structure of bromide-bound *Gt*ACR1. The structure in bromide provides direct evidence for our proposed conductance mechanism (*Li et al., 2019*) and also demonstrates protein conformational changes in C1 and C3, which shed light into the role of the constrictions in channel opening.

## Results and Discussion

### Overall structure of bromide-bound GtACR1

To facilitate incorporation of bromide, 100 mM NaBr was supplied in both protein purification and crystallization buffers. To avoid radiation damage, the *in meso in situ* serial data collection method (IMISX) (*Huang et al., 2018*) was used. The structure of bromide-bound *Gt*ACR1 was determined at 3.2 Å resolution with merging 36 partial data sets by molecular replacement (MR) using the apo form structure (PDB code 6EDQ) as the model (Table 1).

**Table 1.**
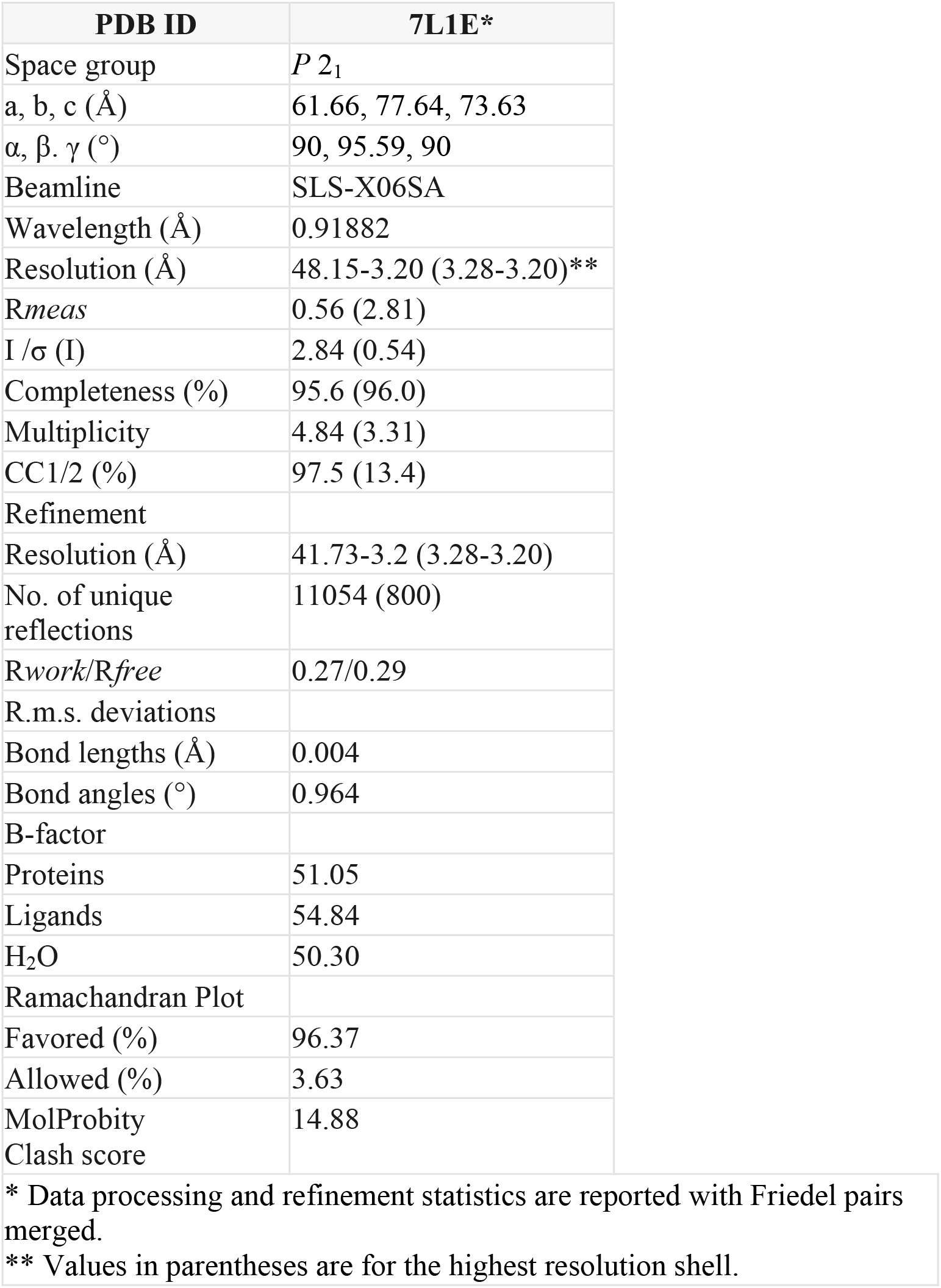
Crystallographic data and refinement of the bromide-bound *Gt*ACR1 structure*.

The bromide-bound *Gt*ACR1 exhibits a similar homodimeric overall structure as the apo form (rmsd: 0.6 Å by comparing Cα from residues 1-295) (Fig. 1A). All 7-helices and trans-configured retinal moieties are well superimposed including the inter-subunit disulfide bridge stabilizing the N-terminus fragments of the two protomers on the extracellular surface. Each protomer exhibits a continuous tunnel extending from the extracellular to intracellular surfaces of the protein similar to that seen in the apo form structure (Fig. 1A and 1B).

**Figure 1.**
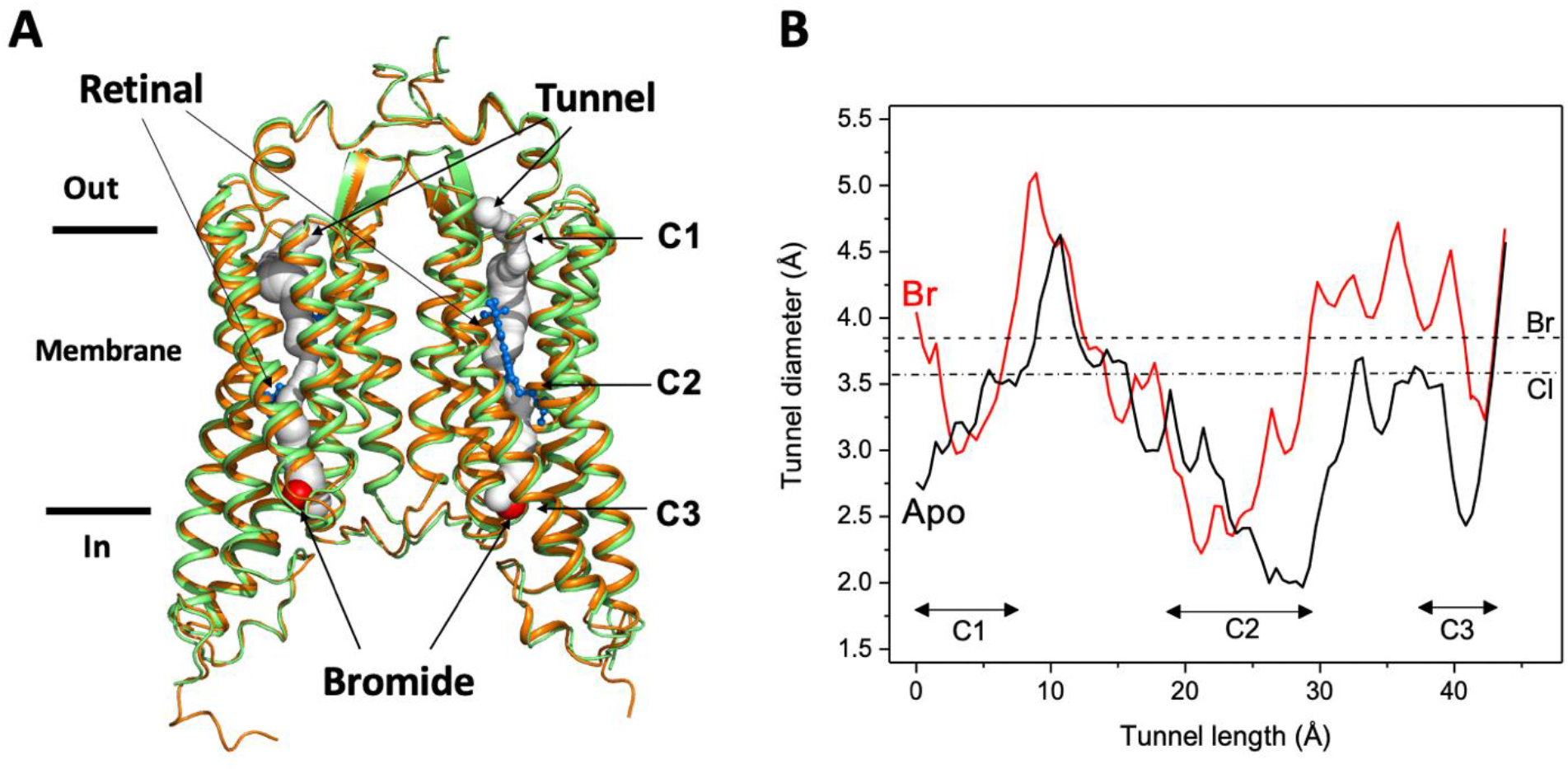
Overall conformation of the bromide-bound *Gt*ACR1 structure. **A:** Superimposition of *Gt*ACR1 apo (*orange*, PDB 6EDQ) and bromide-bound (*green*) structures; one bromide ion (*red* sphere) is located at the cytoplasmic entry of the intramembrane tunnel (*grey* tube, predicted using the program CAVER (*Chovancova et al., 2012*) of each promoter. All-trans-retinal moieties are depicted as *blue* sticks. **B:** Tunnel profile of *Gt*ACR1 protomer B predicted by CAVER: *Gt*ACR1 apo form (*black* line); bromide-bound form (*red* line). The sizes of chloride and bromide ions are indicated as dot-dashed and dashed lines, respectively.

As seen from comparison of the tunnel profile diameters of the apo and bromide bound structures (Fig. 1B) bromide binding enlarges the tunnel in constriction C3 on the cytoplasmic side of the tunnel, where it is bound (Fig. 1A), and also in part of C2, adjacent to the binding site, and the more distant constriction C1 on the extracelluar side of the tunnel. The widened constriction regions indicate a preactivated conformation exhibiting a closed channel with fractional transition to an open-channel state.

### Bromide binding at the tunnel entry

A bromide ion was found at the cytoplasmic port of the tunnel in each protomer (Fig. 1A & Suppl. Fig. 1A). *A priori*, given the larger number of electrons in Br (Z=35), it would be difficult to mistake it for a water molecule (Z=10), but to test this possibility directly, the bromide ion was replaced with a water molecule and the structure refined using *PHENIX* (*Adams et al., 2010*). The refinement showed a strong positive electron density at the bromide position in the *Fo-Fc* difference map and it was diminished only when a bromide ion was placed at that position (Suppl. Fig. 1B). This evidence excludes a water molecule as responsible for the electron density at the position. We conclude a bromide ion resides at the tunnel entry in each protomer.

The bromide binding site is formed by three cyclic residues: Pro58 from the cytoplasmic loop between TM1 and 2, and Trp246 and Trp250 from TM7 in a triangular configuration (Fig. 2A). In the binding site, a bromide is stabilized by a H-bond interaction formed by the indole NH group of Trp246. This type of anion binding conformation has also been found in several chloride-bound nucleotide structures (*Auffinger et al., 2004*). Pro58 may play an important role in substrate binding by pressing its ring towards the bromide anion with a short distance of 2.4 Å. Unlike other aromatic residues, the ring of proline exhibits a partial positive charge due to electron withdrawal by the adjacent protein backbone and the lower electronegativity of the hydrogens on the ring surface (*Zondlo, 2013*). The partial electropositivity of Pro58 may contribute to the binding of bromide via electrostatic interactions. Notably, such a close proline-halide interaction has also been observed in the structure of the chloride-pump rhodopsin ClR (site 2) (*Kim et al., 2016*).

**Figure 2.**
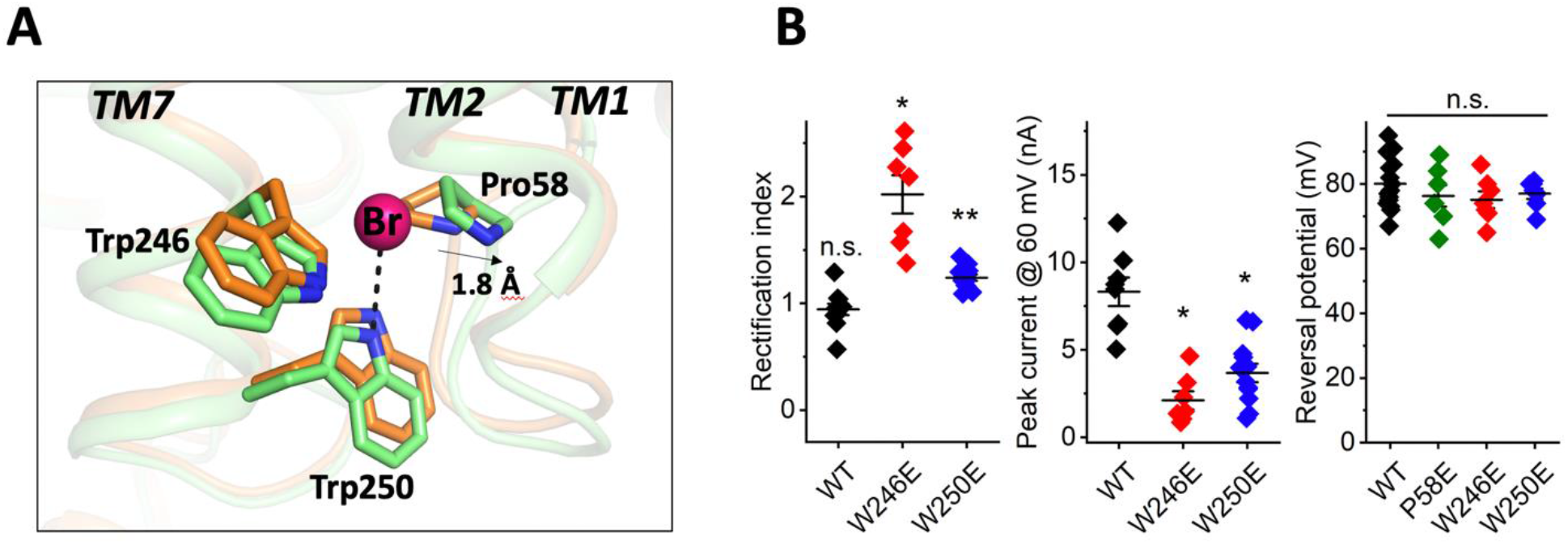
Structure of the bromide binding site in the apo and bromide-bound *Gt*ACR1 and electrophysiological properties of site mutants. **A:** A bromide ion (*red* sphere) stabilized by three cyclic residues (*green* sticks) via H-bond interaction (*black* dashed line) with superimposition of the apo form structure (*orange*). **B:** Functional probing of the bromide binding site residues by patch clamp analysis of their mutants: *Left*: Rectification index (RI), defined as the ratio of peak photocurrent amplitudes recorded at + 60 and - 60 mV at the amplifier output. RI > 1 by one-sample Wilcoxon signed-rank test: * p < 0.05, ** p < 0.01, n.s. not significant (p > 0.05). *Middle:* Peak current at 60 mV. Comparison with the wild-type by Mann-Whitney test: * p < 0.005. *Right:* Reversal potential at the reduced Cl^-^ concentration in the bath. Comparison with the wild-type by Mann-Whitney test: n.s., not significant (p > 0.05).

Our previous results implicated Pro58 in gating of *Gt*ACR1, in that substitution of Pro58 by Glu reduced photocurrent amplitude, altered the kinetics of channel closing, and caused strong outward rectification of the current-voltage dependence (*Li et al., 2019; Sineshchekov et al., 2019*). We observed similar effects in W246E and W250E mutants in the bromide binding site (Fig. 2B), in that photocurrent amplitudes were significantly reduced compared to the wild type and outward rectification was increased (Fig. 2B). None of these three substitutions with Glu reduced the selectivity for anions, as assessed by reversal potential measurements (Fig. 2B).

### Conformational changes of the C1 and C3 constrictions

Despite the similar overall structure to that of the apo form, conformational changes were observed at the C1 and C3 constrictions within the tunnel. Pro58 is an important component of C3. In the apo form structure, Pro58, together with Leu108, Ala61, and Leu245, constrain the cytoplasmic port, leading to the cytoplasmic half of the tunnel narrowing to 3.6 Å in diameter (*Li et al., 2019*). In the bromide-bound structure, the presence of bromide pushes Pro58 outward by ~2 Å (Fig. 2A). As a result, the cytoplasmic half of the tunnel is broadened by 1Å in diameter (Fig. 1B).

Conformational changes were also observed in the extracellular half of the tunnel. In the apo form structure, the C1 constriction is stabilized by a salt-bridge formed by Arg94 and Glu223 near the extracellular surface (Fig. 3A). In the bromide-bound structure, Arg94 undergoes salt-bridge switching along the tunnel. The side-chain of Arg94 is flipped by ~180° to form an alternative salt-bridge with Asp234 in the photoactive site (Fig. 3A), resulting in modest relaxation (~1 Å in diameter) of the C1 constriction (Fig. 1B).

**Figure 3.**
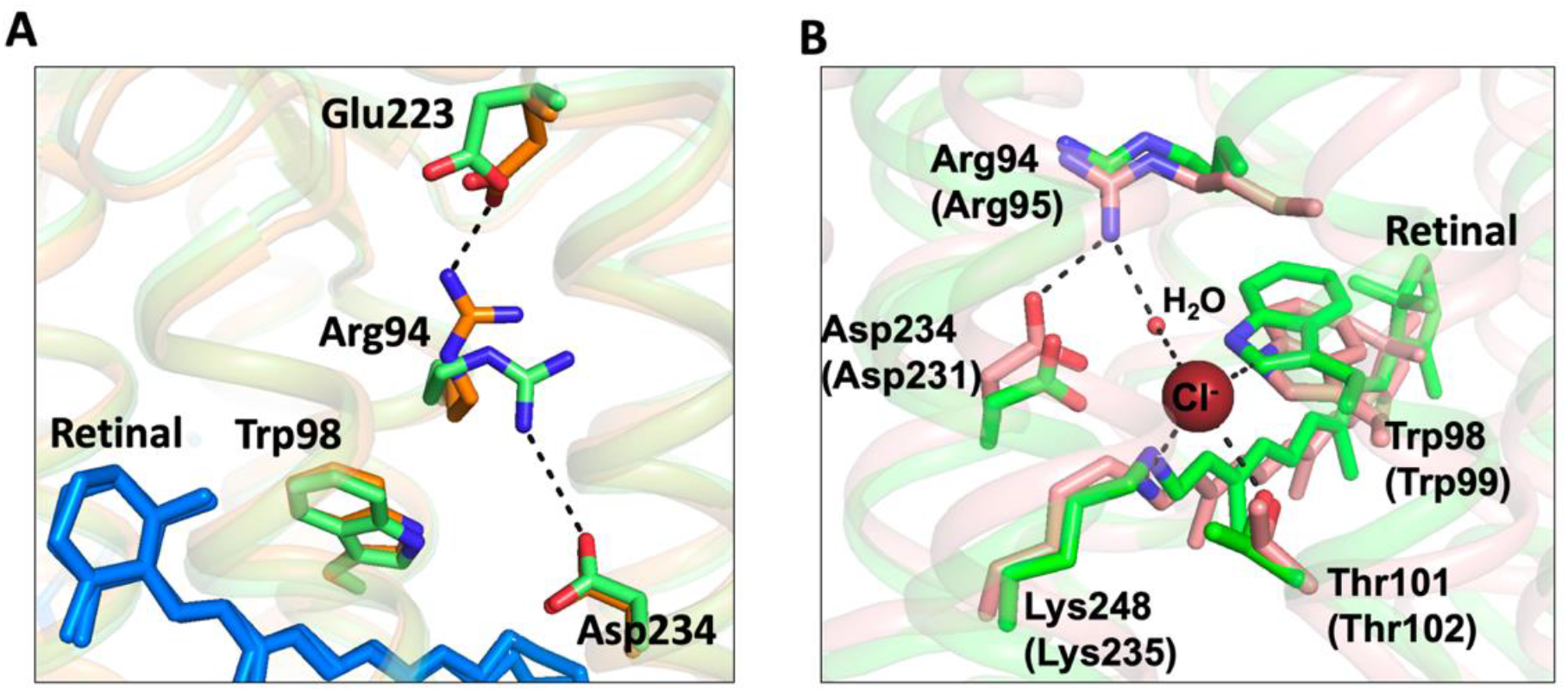
Conformational changes of the C1 constriction. **A:** Superimposition of *Gt*ACR1 apo form (*orange*) and bromide-bound (*green*) structures showing salt-bridge switching (*black* dashed lines) of Arg94 from the extracellular Asp223 to Asp234 near the mid-membrane retinal (*blue* sticks). **B:** Superimposition of the bromide-bound *Gt*ACR1 (*green*) and Cl-pump CIR (*pink*, PDB 5G2A) structures showing a similar halide binding site in the extracellular half of the tunnel. In the CIR structure, a chloride ion (*maroon* sphere) is stabilized via H-bond interactions (*black* dashed lines). The residues are labelled as in *Gt*ACR1 and analogous residues for CIR are indicated in parentheses.

Arg94 is highly conserved in the microbial rhodopsin family and it is critical in maintaining anion conductance of *Gt*ACR1. The mutation R94A nearly abolished anion conductance (*Li et al., 2019*). Arg94 is the only positively-charged residue in the extracellular half of the tunnel. It may enable transfer of anions across the extracellular half of the tunnel via charge-charge interaction. We found that this side chain rotation enables Arg94 and its neighboring residues to form a conformation nearly identical to the chloride binding site of the Cl-pump CIR (*Kim et al., 2016*) (Fig. 3B). In the structure of CIR, a chloride ion (site 1) is bound between Arg95 and the Schiff base via salt-bridges. Although some densities were also observed near Arg94 in bromide-bound *Gt*ACR1, those weak densities prevent unambiguous determination of any halide anion in the vicinity. However, the similar conformations (Fig. 3B) suggest that Arg94 rotates its side chain to form a functional anion binding site with the Schiff base in *Gt*ACR1.

## Conclusion

In this research advance, we addressed several major questions raised by our previous apo form structure by determining the crystal structure of bromide-bound *Gt*ACR1. We identified a novel bromide binding site at the cytoplasmic entry of the transmembrane tunnel (Fig. 1). This finding provides direct evidence for our hypothesis (*Li et al., 2019*) that the tunnel we observe both in the apo form and, in an altered form, in the bromide-bound condition is the closed anion channel. The structure shows protein conformational changes induced by bromide binding that widen the tunnel: (1) bromide binding at the cytoplasmic entry induces relaxation of the C3 constriction (Fig. 2A); and (2) salt-bridge switching of Arg94 may create an anion binding site with the protonated Schiff base (Fig. 3A & 3B). These observations indicate that substrate binding induces a transition from an inactivated state to a pre-activated state in the dark. Despite the conformational changes that expand the tunnel in C1 and C3, the narrowest constriction C2 remains tightly closed (Fig. 1B), which is attributable to the all-*trans*-configured retinal moiety within the tunnel. These results suggest a dominant role of the photoactive site Schiff base *per se* in the channel-gating mechanism. Moreover, the pre-activating conformational changes induced by bromide binding may facilitate channel opening by reducing free energy in the tunnel constrictions, consistent with the larger conductance of bromide vs chloride (*Govorunova et al., 2015*). Future structural study of *Gt*ACR1 in light-activated states is needed to resolve protein conformational changes in its photochemical reaction cycle.

## Methods

### Protein expression and purification

Protein expression and purification of *Gt*ACR1 expressed in *Pichia pastoris* followed the procedure described (*Li et al., 2019*). The eluted protein was further purified using a Superdex Increase 10/300 GL column (GE Healthcare, Chicago, IL) equilibrated with buffer containing 350 mM NaBr, 5% glycerol, 0.03% DDM, 20 mM MES, pH 5.5, thereby replacing Cl^-^ with Br^-^ in the micelle suspension. Protein fractions with an A280/A515 absorbance ratio of ~1.9 were pooled, concentrated to ~20 mg/ml using a 100 K MWCO filter, flash-frozen in liquid nitrogen, and stored at −80°C until use. Molar protein concentration was calculated using the absorbance value at 515 nm divided by the extinction coefficient 45,000 M^-1^cm^-1^.

### Protein crystallization

Crystallization was carried out using the in meso method as with the apo protein (*Li et al*., *2019*). The lipidic mesophase (lipidic cubic phase, LCP) sample was obtained by mixing 40 μl of *Gt*ACR1 protein with 60 μl monoolein (MO) (Sigma, St. Louis, MO or Nu-chek, Waterville, MN) using two syringes until the mixture became transparent. Crystallization trials were setup using both 96-well LCP glass sandwich plates (Molecular Dimensions, Maumee OH) and IMISX™ plates (MiTeGen) which are designed to perform *in meso in situ* serial X-ray crystallography (*Huang et al., 2018; Huang et al., 2016*) with 150 nl aliquots of the protein-mesophase mixture and overlaid with 1.5 μl of precipitant solution using a Gryphon crystallization robot (Art Robbins, Sunnyvale, CA). The plates were covered by aluminum foil to maintain a dark environment and incubated at room temperature. Red-colored *Gt*ACR1 crystals of ~20 μm in size appeared after one month. The most highly diffracting crystals were obtained from crystallization screen containing 15% 2-methyl-2,4-pentanediol (MPD), 0.1 M NaBr and buffer of 0.1 M MES, pH 5.5, or Na-acetate, pH 5.5. LCP crystals from glass plate were harvested using micromesh loops (MiTeGen, Ithaca, NY) and the wells with crystals-laden LCP in IMISX plate were retrieved using a glass cutter and scissors and mounted using 3D-printed holders (*Huang et al., 2020*). All the samples were flash-cooled in liquid nitrogen without any additional cryoprotectant and stored in uni-pucks (MiTeGen, Ithaca, NY) for X-ray diffraction. An improvement over our previous work was setting up the crystallization in the IMISX™ plates and shipping the plates to PSI. This step prevents potential damage to the crystals during harvesting and facilitates high-throughput screening for diffractable crystals.

### Data collection and processing

X-ray diffraction data collections were performed on protein crystallography beamlines X06SA-PXI at the Swiss Light Source (SLS), Villigen, Switzerland. Data were collected with a 10 × 10 μm^2^ micro-focused X-ray beam of 13.49 keV (0.91882 Å in wavelength) at 100 K using SLS data acquisition software suites (DA+) (*Wojdyla et al., 2018*). Continuous grid-scans (*Wojdyla et al., 2018*) were used to locate crystals in frozen LCP samples both from conventional loop and IMISX samples (*Huang et al., 2016*). The crystals harvested on loop were collected by the rotation method with 0.2 s exposure time, 0.2° oscillation for data collection and 30° wedge for each crystals. The sample using IMISX setup were measured by an automated serial data collection protocol (CY+) as described (*Basu et al., 2019*) using the following parameters: 0.2 ° oscillation and 0.1 s exposure time for data collection with 10-20° wedge for each crystal. The EIGER 16M detector operated in continuous/shutterless data collection mode. Data were processed with XDS and scaled and merged with XSCALE (*Kabsch, 2010a; Kabsch, 2010b*). The data sets were further selected to improve the final merging data set with the XDSCC12 (*Assmann et al., 2020*). 36 partial data sets were collected, processed, and merged to a final data set to 3.2 Å resolution, in which 31 data sets were collected on the crystals in the IMISX setup and 5 data sets were collected from crystals harvested on loop. Data collection and processing statistics are provided in Table 1.

### Structure determination and analysis

The structure of bromide-bound *Gt*ACR1 was determined by the molecular replacement (MR) method using 6EDQ (*Li et al*., *2019*) as the search model with the program Phaser (*McCoy et al., 2007*). The structure was refined using PHENIX (*Adams et al., 2010*) and model building was completed manually using COOT (*Emsley and Cowtan, 2004*). The final structure yields a R_work_/R_free_ factor of 0.26/0.29. Refinement statistics are reported in Table 1. The structure factors and coordinates have been deposited in the Protein Data Bank (PDB entry code: 7L1E). Figures of molecular structures were generated with PyMOL (http://www.pymol.org). We analyzed the halide tunnel using the program CAVER with 0.9 nm as the detecting probe (*Chovancova et al., 2012*).

### Electrophysiology of GtACR1 mutants

*Gt*ACR1 mutants were characterized by whole-cell patch clamp recording as described in detail in our previous report (*Li et al., 2019*). Briefly, the wild-type expression construct was cloned into the mammalian expression vector pcDNA3.1 (Life Technologies, Carlsbad, CA) in frame with an EYFP (enhanced yellow fluorescent protein). Mutations were introduced using a QuikChange XL site-directed mutagenesis kit (Agilent Technologies, Santa Clara, CA) and verified by DNA sequencing. HEK293 (human embryonic kidney) cells were transfected using the ScreenFectA transfection reagent (Waco Chemicals USA, Richmond, VA). All-*trans*-retinal (Sigma, St. Louis, MO) was added at the final concentration 4 μM immediately after transfection. Photocurrents were recorded 48-72 h after transfection in whole-cell voltage clamp mode at room temperature (25°C) with an Axopatch 200B amplifier (Molecular Devices, Union City, CA) and digitized with a Digidata 1440A using pClamp 10 software (both from Molecular Devices). Patch pipettes were fabricated from borosilicate glass and filled with the following solution (in mM): KCl 126, MgCl2 2, CaCl2 0.5, EGTA 5, HEPES 25, and pH 7.4. The standard bath solution contained (in mM): NaCl 150, CaCl_2_ 1.8, MgCl_2_ 1, glucose 5, HEPES 10, pH 7.4. To test for changes in the permeability for Cl^-^, this ion in the bath was partially replaced with non-permeable aspartate (the final Cl^-^ concentration 5.6 mM). For each cell, a single value of the Erev was calculated. The holding potential values were corrected for liquid junction potentials calculated using the Clampex built-in LJP calculator. Continuous light pulses were provided by a Polychrome V light source (T.I.L.L. Photonics GMBH, Grafelfing, Germany) at 15 nm half-bandwidth in combination with a mechanical shutter (Uniblitz Model LS6, Vincent Associates, Rochester, NY; half-opening time 0.5 ms). The maximal light intensity at the focal plane of the objective lense was 7.7 mW mm^-2^ at 515 nm.

Batches of culture were randomly allocated for transfection with a specific mutant; no masking (blinding) was used. Individual transfected HEK293 cells were selected for patching by inspecting their tag fluorescence; non-fluorescent cells were excluded, as were cells for which we could not establish a gigaohm seal. Results obtained from different individual cells were considered as biological replicates. The raw data obtained in individual cells are shown as diamonds. Sample size was estimated from previous experience and published work on a similar subject, as recommended by the NIH guidelines. No outliers were excluded. Normality of the data was not assumed, and therefore non-parametric statistical tests were used as implemented in OriginPro 2016 software; P values > 0.05 were considered not significant.

### Cell lines

Only a commercially available cell line authenticated by the vendor (HEK293 from ATCC) was used; no cell lines from the list of commonly misidentified cell lines were used. The absence of micoplasma contamination was verified by Visual-PCR mycoplasma detection kit (GM Biosciences, Frederick, MD).

## Acknowledgements

This work was supported by National Institutes of Health Grant R01GM027750 and Endowed Chair AU-0009 from the Robert A. Welch Foundation to JLS, and American Heart Association Grant 18TPA34230046 to LZ. C-YH was partially supported by the European Union’s Horizon 2020 research and innovation programme under the Marie-Skłodowska-Curie grant agreement No. 701647. The authors thank Yumei Wang for her technical assistance and the assistance and support of beamline scientists at the Swiss Light Source beamlines X06SA-PXI.

---

**Supplementary Figure 1.**
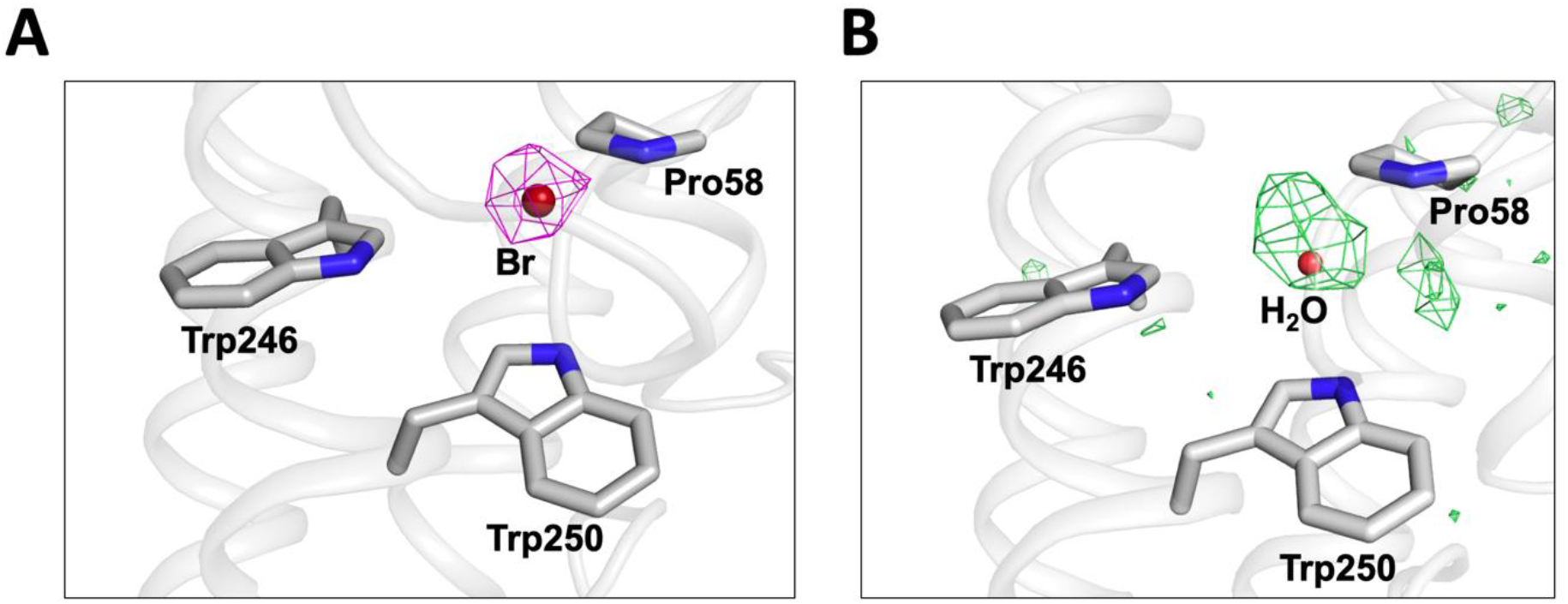
Confirmation of a bromide ion at the cytoplasmic port of *Gt*ACR1. **A:** Bromide ion (*dark red* sphere) in the binding site (*grey* sticks) overlayed with the composite omit map depicted as *magenta* mesh (contoured at 2σ). The composite omit map was calculated using *PHENIX (Adams et al., 2010*). **B:** *F_o_-F_c_* difference map generated by the refinement of a water molecule at the bromide position showing positive electron density depicted as *green* mesh (contoured at +3σ) at the water position.

## References

Adams, P.D., P.V. Afonine, G. Bunkoczi, V.B. Chen, I.W. Davis, N. Echols, J.J. Headd, L.W. Hung, G.J. Kapral, R.W. Grosse-Kunstleve, A.J. McCoy, N.W. Moriarty, R. Oeffner, R.J. Read, D.C. Richardson, J.S. Richardson, T.C. Terwilliger, and P.H. Zwart. 2010. PHENIX: a comprehensive Python-based system for macromolecular structure solution. Acta Crystallogr D Biol Crystallogr. 66:213–221.

Assmann, G.M., M. Wang, and K. Diederichs. 2020. Making a difference in multi-data-set crystallography: simple and deterministic data-scaling/selection methods. Acta Crystallogr D Struct Biol. 76:636–652.

Auffinger, P., L. Bielecki, and E. Westhof. 2004. Anion Binding to Nucleic Acids. Structure (London, England: 1993). 12:379–388.

Basu, S., J.W. Kaminski, E. Panepucci, C.Y. Huang, R. Warshamanage, M. Wang, and J.A. Wojdyla. 2019. Automated data collection and real-time data analysis suite for serial synchrotron crystallography. J Synchrotron Radiat. 26:244–252.

Chovancova, E., A. Pavelka, P. Benes, O. Strnad, J. Brezovsky, B. Kozlikova, A. Gora, V. Sustr, M. Klvana, P. Medek, L. Biedermannova, J. Sochor, and J. Damborsky. 2012. CAVER 3.0: a tool for the analysis of transport pathways in dynamic protein structures. PLoS Comput Biol. 8:e1002708.

Emsley, P., and K. Cowtan. 2004. Coot: model-building tools for molecular graphics. Acta Crystallogr D Biol Crystallogr. 60:2126–2132.

Govorunova, E.G., O.A. Sineshchekov, R. Janz, X. Liu, and J.L. Spudich. 2015. Natural light-gated anion channels: A family of microbial rhodopsins for advanced optogenetics. Science. 349:647–650.

Huang, C.Y., N. Meier, M. Caffrey, M. Wang, and V. Olieric. 2020. 3D-printed holders for in meso in situ fixed-target serial X-ray crystallography. J Appl Crystallogr. 53:854–859.

Huang, C.Y., V. Olieric, N. Howe, R. Warshamanage, T. Weinert, E. Panepucci, L. Vogeley, S. Basu, K. Diederichs, M. Caffrey, and M. Wang. 2018. In situ serial crystallography for rapid de novo membrane protein structure determination. Commun Biol. 1:124.

Huang, C.Y., V. Olieric, P. Ma, N. Howe, L. Vogeley, X. Liu, R. Warshamanage, T. Weinert, E. Panepucci, B. Kobilka, K. Diederichs, M. Wang, and M. Caffrey. 2016. In meso in situ serial X-ray crystallography of soluble and membrane proteins at cryogenic temperatures. Acta crystallographica. Section D, Structural biology. 72:93–112.

Kabsch, W. 2010a. Integration, scaling, space-group assignment and post-refinement. Acta Crystallogr D Biol Crystallogr. 66:133–144.

Kabsch, W. 2010b. Xds. Acta Crystallogr D Biol Crystallogr. 66:125–132.

Kim, K., S.K. Kwon, S.H. Jun, J.S. Cha, H. Kim, W. Lee, J.F. Kim, and H.S. Cho. 2016. Crystal structure and functional characterization of a light-driven chloride pump having an NTQ motif. Nat Commun. 7:12677.

Kim, Y.S., H.E. Kato, K. Yamashita, S. Ito, K. Inoue, C. Ramakrishnan, L.E. Fenno, K.E. Evans, J.M. Paggi, R.O. Dror, H. Kandori, B.K. Kobilka, and K. Deisseroth. 2018. Crystal structure of the natural anion-conducting channelrhodopsin GtACR1. Nature. 561:343–348.

Li, H., C.-Y. Huang, E.G. Govorunova, C.T. Schafer, O.A. Sineshchekov, M. Wang, L. Zheng, and J.L. Spudich. 2019. Crystal structure of a natural light-gated anion channelrhodopsin. eLife. 8:e41741.

McCoy, A.J., R.W. Grosse-Kunstleve, P.D. Adams, M.D. Winn, L.C. Storoni, and R.J. Read. 2007. Phaser crystallographic software. J Appl Crystallogr. 40:658–674.

Sineshchekov, O.A., E.G. Govorunova, H. Li, X. Wang, and J.L. Spudich. 2019. Opposite Charge Movements Within the Photoactive Site Modulate Two-Step Channel Closing in GtACR1. Biophys J. 117:2034–2040.

Wojdyla, J.A., J.W. Kaminski, E. Panepucci, S. Ebner, X. Wang, J. Gabadinho, and M. Wang. 2018. DA+ data acquisition and analysis software at the Swiss Light Source macromolecular crystallography beamlines. J Synchrotron Radiat. 25:293–303.

Zondlo, N.J. 2013. Aromatic-proline interactions: electronically tunable CH/π interactions. Acc Chem Res. 46:1039–1049.

